# Larval Habitat Bacteria Emit Volatile Semiochemicals that Modulate Oviposition Site Selection in the Malaria Vector *Anopheles gambiae* (Diptera: Culicidae)

**DOI:** 10.64898/2026.07.29.741650

**Authors:** Josphat Mutinda, Kennedy Omondi Oduor, Samuel Mwakisha Mwamburi, Maurice O. Omolo, Regina Mongina Ntabo, James Muhunyu Gathiru, Joseph Mwangangi, James Nonoh

## Abstract

**Background:** Larval habitat management is increasingly recognized as a complementary strategy for malaria vector control. While bacterial communities are abundant in mosquito breeding sites, their role in shaping oviposition behavior of *Anopheles gambiae* (Diptera: Culicidae) remains poorly understood. This study evaluated the relative attractiveness or repellency of water infused with bacterial isolates obtained from mosquito larval habitats to gravid *Anopheles gambiae*.

**Methods:** We isolated and taxonomically characterized bacteria from 30 larval habitats in coastal Kenya using 16S rRNA sequencing. Thirty-two representative isolates were tested in two-choice oviposition bioassays with gravid *An. gambiae s.s.*, and volatile organic compounds (VOCs) emitted were analyzed by GC–MS. Statistical analyses included Student’s t-tests, Mann–Whitney U tests, and calculation of the Oviposition Activity Index (OAI).

**Results:** Proteobacteria dominated larval habitats (78%), with frequent genera including *Aeromonas*, *Acinetobacter*, *Pseudomonas*, *Enterobacter*, and *Cronobacter*. Most isolates exerted strong deterrent effects, with *Aeromonas hydrophila*, *Enterobacter hormaechei*, and *Pseudomonas mendocina* completely inhibiting oviposition (OAI = −1; 100% repellency). Other isolates such as *Neobacillus drentensis* and *Aeromonas veronii* reduced egg deposition by >85%. In contrast, a small subset (*Aquitalea pelogenes*, *Pseudomonas oleovorans*, *Cronobacter sakazakii*) showed weak, non-significant attraction (OAI = 0.15–0.23). GC–MS revealed that highly repellent isolates emitted abundant benzenoids, ketones, phenols, ethers, oxygenated heterocycles, and aldehydes.

**Conclusions:** Bacterial volatiles strongly modulate oviposition site selection in *An. gambiae*, with most isolates acting as potent repellents. These findings highlight microbial VOCs as promising candidates for novel vector control strategies targeting gravid mosquitoes. Limitations include the laboratory-based bioassay design and lack of field validation of VOC activity. Future work should identify specific bioactive compounds and evaluate their efficacy under natural conditions.

**Research Highlights:**

- Bacteria from mosquito larval habitats influence oviposition of *Anopheles gambiae*.
- Several bacterial isolates significantly repel gravid mosquitoes.
- GC–MS analysis revealed VOCs including benzenoids, ketones, phenols, and aldehydes.
- Bacterial volatiles show potential for novel mosquito control strategies.

## INTRODUCTION

Malaria is a serious disease caused by *Plasmodium* parasites, which are mainly transmitted by the highly anthropophilic, endophilic, and endophagic female *Anopheles gambiae* in Sub-Saharan Africa (1,2). Although there has been a significant reduction in malaria transmission in most parts of Kenya, the transmission rates remain considerably high among the inhabitants of the coastal region of Kenya (3,4). Intense transmission has been reported to occur in villages near large water bodies, such as lakes and oceans (5). Some available evidence suggests mosquito development does not occur within the water body itself but rather along the shores of these water bodies on some artificial sites such as animal hoofprints and other shallow waters (6). Nevertheless, there is scanty information to describe the ecology of mosquito development sites.

To date, vector control remains the most effective way to prevent malaria. Most vector control strategies have relied on chemical insecticides, targeting the indoor host-seeking behavior of mosquitoes (7). Despite remarkable success, the elimination of malaria remains a big challenge since the malaria parasite is maintained by mosquitoes, which oviposit, feed, and rest in the outdoor environment (8). Because of this and the emergence of insecticide-resistant mosquito vectors and highly drug-resistant malaria parasites (9), there is an urgent need to focus on the larval habitat management and control of oviposition sites seeking malaria vectors (10). It is believed that targeting the larval stage in malaria vector control is a potentially important tool for malaria eradication. This can be achieved through the modification of larval habitats to make them unsuitable for mosquito larvae to survive (11).

Reproductive fitness, coupled with the availability of adequate and suitable proliferation sites, is believed to be a major factor influencing the abundance and distribution of mosquito species (12). The choice of appropriate oviposition sites by gravid mosquitoes is determined by different physical, chemical, and environmental cues such as attractants, repellents, or stimulants (13). Oviposition site search and selection is a very dynamic process where the gravid mosquito integrates several environmental signals, such as humidity, olfactory, tactile, thermal, and visual cues to assess the suitability of oviposition sites (13). Chemicals mediating oviposition in mosquitoes have been identified from sources like hay infusion, water-associated bacteria, larvae holding water, pheromones, and exudates from aquatic competitors or predators. These volatile semiochemicals emitted from the mosquito oviposition sites are important as initial interspecific attractants (14). Furthermore, microbial populations in mosquito proliferation sites produce metabolites that are associated with positive oviposition behavior (14).

The size of the adult mosquito population is largely dependent on bacteria present in the mosquito larval habitats (15). Bacteria form part of the biotic factors in mosquito larval habitats and therefore play a very significant role in the life cycle of mosquitoes. Bacteria are the most important microbial constituents of mosquito larvae’ food, and published data show that mosquitoes can grow on cultures made only of bacteria (16). A direct oviposition response by mosquitoes toward bacterial cultures or filtrates has been reported. These bacteria have also been reported as direct sources of food for the larvae (18). Bacteria such as *Pseudomonas, Bacillus, Enterobacter, Klebsiella, Aeromonas, and Acinetobacter* are widely distributed and abundant in mosquito breeding sites and have been suggested as important sources of food for the larva (11). Bacteria in mosquito proliferation sites have also been discovered to affect mosquito susceptibility to various pathogens and the resistance of mosquitoes to pesticides (19). Moreover, it has been reported that these bacteria significantly influence mosquito oviposition behavior (20).

Volatile semiochemicals of microbial origin serve as attractants or stimulants to different species of oviposition-seeking gravid mosquitoes (14). It has been reported that oviposition site selection and preference by *Anopheles* mosquitoes can be influenced by semiochemicals of microbial origin (12). Gravid *Anopheles gambiae* preferred to lay more eggs on non-autoclaved substrates taken from natural larval habitats than on similar but autoclaved materials (14), potentially suggesting that destroying the microorganisms by autoclaving eliminated the source of semiochemicals that made these substrates attractive to the gravid mosquitoes.

The origin of oviposition stimulants in mosquito proliferation sites and their mode of action towards gravid females have been suggestively linked to microbial activity, though not fully understood (20). Bacteria in larval habitats help in the modification of organic matter in mosquito breeding sites, which may give rise to chemical constituents that are ingested by larvae as well as volatile organic compounds mediating the oviposition site selection behavior of mosquitoes (16). Such bacteria may play important roles in the production of different volatile blends that influence malaria mosquitoes to their preferred oviposition sites, although there is limited information on the profiles of specific bacteria involved in the process as well as the specific volatiles they emit. This study aimed to characterize bacteria isolated from the mosquito breeding sites within Lunga Lunga Sub-County in coastal Kenya, evaluate their effect on the oviposition site selection responses of gravid *Anopheles gambiae* ss and analyse the profiles of the volatile compounds they emit.

## MATERIALS AND METHODS

### Sample Collection

Water samples were collected from mosquito breeding sites in Lunga Lunga Sub-County, Kwale County, Kenya. Sampling targeted accessible sites with or without mosquito larvae. GPS coordinates were recorded using a Garmin GPSMAP 64 (Garmin International Inc., Switzerland) and later used to map the sampling area. Each site was visually inspected, and if larvae were absent, at least ten dips were taken using a standard 350-milliliter dipper (BioQuip products, Rancho Dominguez, USA). Sites were classified as positive if at least one larva was found and negative if none were detected. A total of 30 sites (16 positive and 14 negative) were sampled between June and December 2021, between 7 a.m. to 6 p.m. Sampling covered diverse habitats, including drainage pits, swamps, animal hoofprints, and roadside pools, to ensure representative coverage. From each site, one 500 mL sterile plastic bottles were used to collect water samples for bacterial isolation and characterization. Bottles were rinsed with site water before filling, then sealed and transported in a cooler to the laboratory for analysis.

### Isolation and Characterization of Bacteria

A loopful of each water sample was inoculated onto trypticase soy agar (TSA), eosin methylene blue agar (EMB), nutrient agar (NA), and Reasoner’s 2 agar (R2A) plates and incubated aerobically at 37°C for 24 hours to isolate bacterial colonies. DNA was extracted from bacterial isolates using the CTAB method (21). The 16S rRNA gene was amplified by PCR using universal primers 27f (5’-AGAGTTTGATCCTGGCTCAG-3’) and 1492R (5’- GGTTACCTTGTTACGACTT-3’) (22). Purity and concentration of PCR products were verified by gel electrophoresis and NanoDrop spectrophotometry (23,24). Sequencing of PCR products in forward and reverse directions was performed at Inqaba Biotechnical Industries (Pty) Ltd., South Africa. The 16S rRNA gene sequences were visualized and edited using FinchTV software version 1.4.0 (Geospiza, Inc.). After editing, sequences were exported as FASTA files, and bacterial identities were determined using the BLAST protocol (Gapped BLAST).

### Mosquitoes for Oviposition Bioassays

Fresh *Anopheles gambiae* ss eggs were obtained from the Kenya Medical Research Iinstitute (KEMRI) insectary in Kilifi and transported to the KEMRI laboratory in Kwale under wet conditions at ambient temperature. Upon arrival, eggs were placed in shallow plastic trays with tap water at ambient temperature to hatch. Hatched larvae were distributed across multiple trays to minimize overcrowding and ensure optimal growth. They were fed daily with a pinch of powdered Tetramin baby fish food, and water in the trays was changed every two days to maintain a clean environment. Rearing was conducted under ambient conditions (28 °C, 76.49% humidity, 12:12 light cycles). Pupae were transferred into 250 ml plastic cups using 5 ml droppers and placed in mosquito-rearing cages. The cups were covered with fine cotton nets secured by rubber bands, with a small opening sealed with cotton wool to allow adult mosquito collection while preventing escape.

### Mosquito Rearing Cages

The mosquito-rearing cages were cuboid frames measuring 30 cm × 30 cm × 30 cm, constructed from 6 mm stainless steel rods with a base made of 28-gauge galvanized iron sheets. The cages were covered with fine cotton netting, designed with a sleeve opening for introducing pupae cups and aspirating adult mosquitoes. Adult mosquitoes were fed *ad libitum* with 10% glucose solution soaked in cotton wool, placed on top of the cage to provide nourishment and hydration. After 5–6 days of growth, depending on the ambient temperature, the adult mosquitoes were ready for bioassays.

### Preparation of Mosquitoes for Oviposition Bioassay

Mosquito preparation followed standard methods with minimal adjustments (25,26). A group of 100 female and 100 male 6-day-old *Anopheles gambiae ss* mosquitoes were placed in a standard mosquito cage, maintaining a relative humidity of 75% to 85% by covering the cage with a cotton towel soaked in tap water. After starving the mosquitoes of glucose for 6 hours, they were allowed to blood-feed on a human arm for 15 minutes in total darkness. Mosquitoes that had not been fed were separated into another cage. After the first blood meal, the sugar solution was replaced for 15 hours before the mosquitoes were sugar-starved again for a second blood meal. Following the second feeding, glucose was provided again, and the mosquitoes were kept at ambient temperature and humidity for 72 hours to digest the blood meal and mature eggs. Fully gravid females were selected based on abdominal inspection and transferred in groups of five to oviposition cages.

### Oviposition Bioassay with Bacterial Isolated from the Sites

Cage bioassays were conducted under ambient laboratory conditions using standard mosquito cages, following a modified two-choice, single-cup oviposition method (14). The oviposition cups were 100-ml stainless steel, 5 cm in diameter and 8 cm in height. They were cleaned and conditioned in an oven at 50°C for 3 hours to remove residual volatiles. A filter paper cone was placed on top of each cup, ensuring the tip dipped into the test or control water. Pure bacterial isolates were cultured in 1 ml of tryptic soy broth at 37°C for 24 hours to assess their impact on mosquito oviposition behavior. In the treatment cup, 1 ml of the bacterial suspension in tryptic soy broth was added, followed by 70 ml of sterilized distilled water. A filter paper cone was placed on the cup, its tip just touching the water surface. The control cup was prepared similarly but with sterile tryptic soy broth instead of the bacterial suspension. Both cups were placed diagonally in opposite corners of the oviposition cages containing gravid mosquitoes, and the setup was left overnight. The experiment was repeated five times with different sets of five gravid female *Anopheles* mosquitoes (14). The choice of five replicates was based on a priori power analysis using expected effect sizes from similar oviposition studies (14, 25). With an anticipated large effect size (Cohen’s d ≥ 0.8), five replicates provided >80% statistical power at α = 0.05 to detect significant differences between treatment and control groups. This replication level also aligns with WHO guidelines for mosquito bioassays, ensuring reproducibility while balancing ethical considerations in mosquito use. The following morning, the cups were removed, and the total number of eggs laid on each filter paper was recorded.

### Volatiles Collection and Analysis from Bacterial Isolates

Volatiles from bacterial isolates were collected using 250-ml steel tins (7.4 cm diameter, 7.6 cm height), each equipped with two openings—one for the inlet and one for the outlet—fitted with stainless steel tubes (6 mm internal diameter, 5 cm length). Volatile collection traps packed with Porapak Q adsorbent (80–100 mesh size) were inserted into these tubes. The trap was suspended in the headspace above the substrate during collection. The volatile collection traps were made from 8 cm long, 4 mm diameter glass tubes filled with 20 mg of Porapak Q, with 1 cm of space left empty at both ends. Glass wool was placed at both ends to keep the adsorbent in place. After packing, the traps were cleaned in a Soxhlet apparatus for 7 days with dichloromethane and reconditioned in a 120°C oven for 5 days to expel the solvent.

Conditioned traps were connected to an air entrainment kit via Teflon tubes. The Teflon seal ensured an airtight connection, as Teflon is inert and does not emit volatiles. Clean cotton gloves were used to handle the traps. The air entrainment kit created an airflow through the sample’s headspace to collect volatiles in the traps. For volatile collection, a colony was incubated in 1 ml of trypticase soy broth at 37°C for 24 hours. The bacterial suspension was transferred to a volatile collection cup and diluted with 70 ml of sterilized distilled water. The volatile collection trap was introduced into the headspace through the outlet opening. Volatiles were collected dynamically for 2 hours at a flow rate of 0.6 L/min. After collection, the trap was removed, labeled, and sealed in a glass tube for GC-MS analysis. A control sample with sterile broth was also collected to assess background volatiles.

### GC-MS (Gas Chromatography-Mass Spectrometry) Analysis

The volatiles trapped in the adsorbent were eluted with dichloromethane on ice and transferred into borosilicate glass tubes with Teflon stoppers. The tubes were clearly labeled and stored at - 20°C until analysis. The analysis was performed using an HP GC 1800 II (Agilent Technologies, USA) equipped with a DB-5 MS column (30 m x 0.25 mm, 0.25 µm film thickness). Mass spectra were acquired in EI mode (70 eV) with a scanning time of 1.5 seconds across the m/z range of 0-400 a.m.u. Helium was used as the carrier gas at a flow rate of 1 ml/min and a split ratio of 1:30. The injection temperature was set at 250°C, the detector temperature at 270°C, and the column temperature was programmed from 40°C to 240°C at a rate of 5°C per minute. The volatile chemical constituents were identified based on their retention time (rt) and fragmentation patterns in mass spectra using the NIST and Wiley MS libraries. The concentration of each component was determined by comparing the peak area to that of tridecane, which was used as a reference standard in each test (27).

### Data Analysis

Statistical analyses were performed using XLSTAT (XLSTAT, 2007) and R software (version 4.1.3, R Development Core Team, 2021). The data from the bioassays were first processed to calculate the mean and standard deviation for the number of eggs laid in both the treatment and control groups, based on five replicates. Prior to inferential analysis, data were assessed for normality using the Shapiro–Wilk test and homogeneity of variances using Levene’s test. Where assumptions of normality and homoscedasticity were satisfied, differences in egg counts between treatment and control groups were evaluated using a two-tailed Student’s t-test. In cases where assumptions were violated, the non-parametric Mann–Whitney U test was applied (28). A *two-tailed Z-test* was employed to compare the percentages in different groups to determine if differences between groups were statistically significant. The Oviposition Activity Index (OAI) was calculated to assess mosquito oviposition preference. To quantify oviposition preference, the Oviposition Activity Index (OAI) was calculated as:

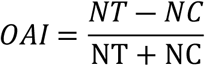

where *NT* represents the number of eggs laid in the test cup and *NC* the number of eggs laid in the control cup. OAI values range from -1 to +1, with values close to 0 indicating no preference, values greater than +0.3 indicating attraction, and values less than -0.3 indicating repellence (29). The OAI was multiplied by 100 to obtain percentage equivalents of the oviposition response for each treatment. The mean and standard deviation of the OAI for each treatment were calculated across the five replicates. Effect sizes (Cohen’s *d*) were calculated for comparisons between treatment and control groups to assess the magnitude of observed differences. Statistical significance was set at p < 0.05. For the GC-MS data analysis, volatiles were identified and quantified based on retention time and mass spectra, utilizing the NIST and Wiley databases. The data were first cleaned by excluding volatiles present in both the sample and control. To examine the relationships between bacterial isolates and the volatile compounds emitted, Principal Component Analysis (PCA) and Partial Least Squares Discriminant Analysis (PLS-DA) were conducted in R.

### Research Permit and Ethical Considerations

Ethical clearance for this study was obtained from the Maseno University Ethics Review Committee (Reference Number: MSU/DRPI/MUERC/803/19). Verbal informed consent was obtained from landowners prior to environmental sampling, as no personal or sensitive data were collected and no procedures posed risk to participants. Written informed consent was also obtained from the insectary technician involved in arm-feeding bioassays, which were conducted under controlled laboratory conditions. A research permit was granted by the National Council for Science, Technology and Innovation (Licence No. NACOSTI/P/21/10048). All procedures complied with relevant institutional and national guidelines for the ethical use of animals and human participants in research.

## RESULTS

### Taxonomic Composition of Bacteria Isolated from Mosquito Larval Habitats

A total of 42 bacterial isolates obtained from mosquito larval habitats; 20 from positive sites and 22 from negative sites, were taxonomically classified into three major phyla: Proteobacteria, Firmicutes, and Deinococcus–Thermus (Supplementary table 1). The bacterial community was strongly dominated by Proteobacteria, which accounted for approximately 78% of all isolates, followed by Firmicutes (18%) and Deinococcus–Thermus (2%). Within Proteobacteria, members of the families Aeromonadaceae, Moraxellaceae, Pseudomonadaceae, and Enterobacteriaceae were most abundant. The family Aeromonadaceae was the dominant group and was represented exclusively by the genus *Aeromonas*, including *Aeromonas sp.*, *A. caviae*, and *A. dhakensis*. The family Moraxellaceae was represented by several species of *Acinetobacter*, including *A. calcoaceticus*, *A. baumannii*, *A. junii*, and *Acinetobacter sp.*, making it one of the most frequently recovered genera. Members of the family Pseudomonadaceae were also common and included *Pseudomonas mendocina*, *P. oleovorans*, *P. alcaligenes*, and *Pseudomonas sp.*The family Enterobacteriaceae comprised *Enterobacter hormaechei*, *Cronobacter sakazakii*, *Klebsiella pneumoniae*, and *Klebsiella sp.*, indicating a diverse assemblage of enteric bacteria in larval habitats. Additional proteobacterial families included Chromobacteriaceae, represented by *Vogesella amnigena* and *Aquitalea pelogenes*; Comamonadaceae, represented by *Acidovorax sp.* and *Hydrogenophaga sp.*; and Rhodobacteraceae, represented by *Thioclava sp.* Firmicutes were dominated by the family Bacillaceae, which included *Bacillus sp.*, *Bacillus niacini*, *Neobacillus drentensis*, and *Priestia megaterium*. Two additional families were recovered at lower frequency: Staphylococcaceae, represented by *Staphylococcus arlettae*, and Planococcaceae, represented by *Jeotgalibacillus marinus*. The phylum Deinococcus–Thermus was represented by a single isolate belonging to the family Deinococcaceae (*Deinococcus sp.*). Overall, the bacterial community associated with mosquito larval habitats was characterized by a high dominance of Proteobacteria, particularly members of the genera *Aeromonas, Acinetobacter, Pseudomonas, Enterobacter, and Cronobacter* (Figure 1).

**Figure 1.**
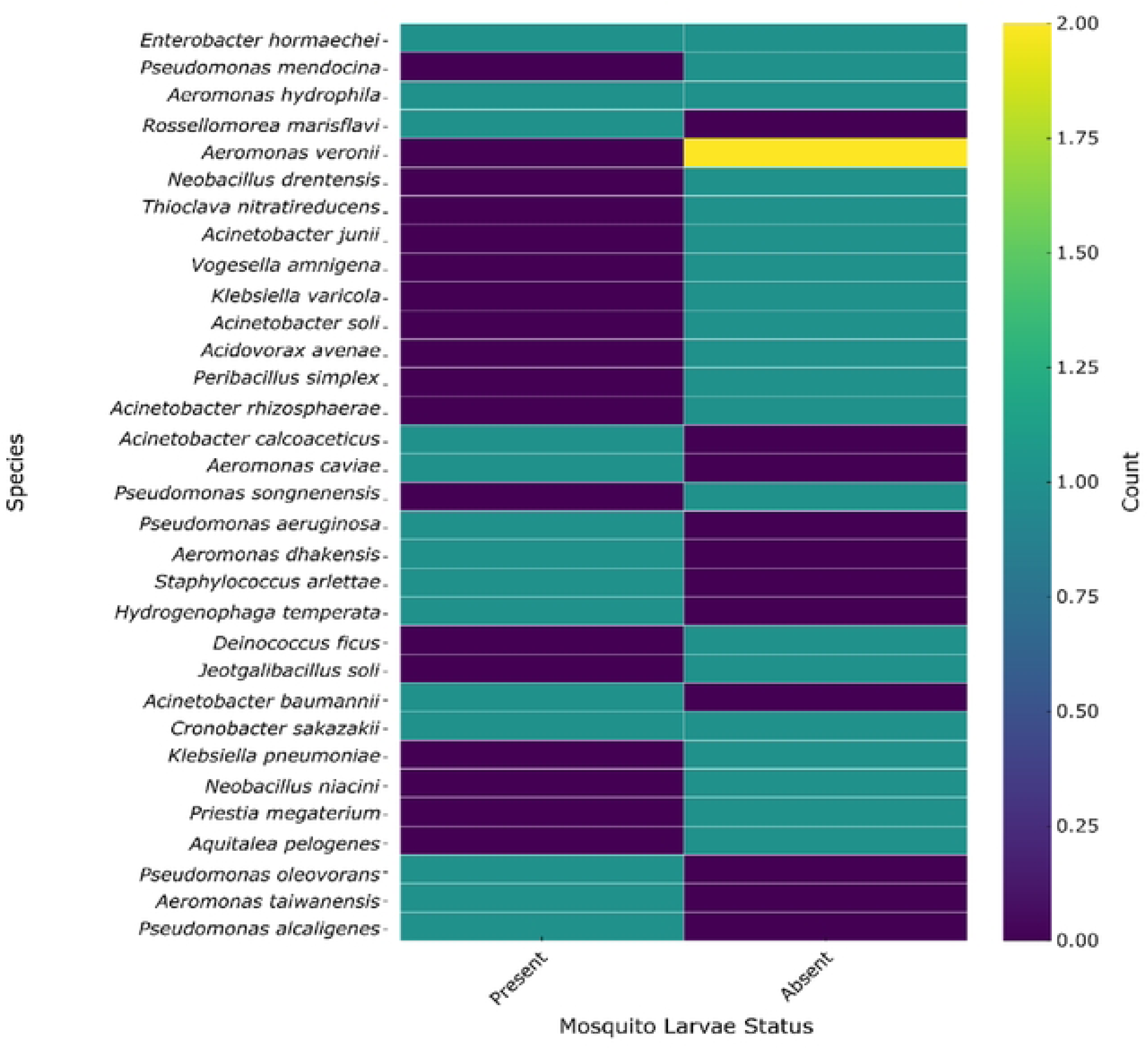
Heatmap showing the distribution of bacterial species in relation to the presence or absence of mosquito larvae. Each row represents a bacterial species, and columns indicate mosquito larvae status. Color intensity corresponds to the count of isolates detected, with darker colors representing higher counts. This visualization highlights patterns of bacterial association with mosquito larvae, suggesting potential microbial indicators or contributors to larval habitat selection and providing insight into microbe–mosquito ecological interactions.

### Oviposition Activity of Bacteria Isolated from Mosquito Larval Habitats

A total of 32 bacterial isolates were tested for their influence on oviposition site selection by *Anopheles gambiae s.s.*. Oviposition bioassays revealed that the majority of bacterial isolates from mosquito larval habitats exerted strong deterrent effects on gravid mosquitoes (Table 1). Several isolates, including *Aeromonas hydrophyla*, *Enterobacter hormaechei*, and *Pseudomonas mendocina*, completely inhibited oviposition, with zero eggs laid in treatment cups compared to 280–531 eggs in controls (P < 0.001), corresponding to an oviposition activity index (OAI) of −1 and 100% effective repellency (ER). Similarly, *Bacillus marisflavi* and *Aeromonas veronii* showed near-complete deterrence, reducing egg deposition by 99.1% and 89.2%, respectively.

**Table 1.**
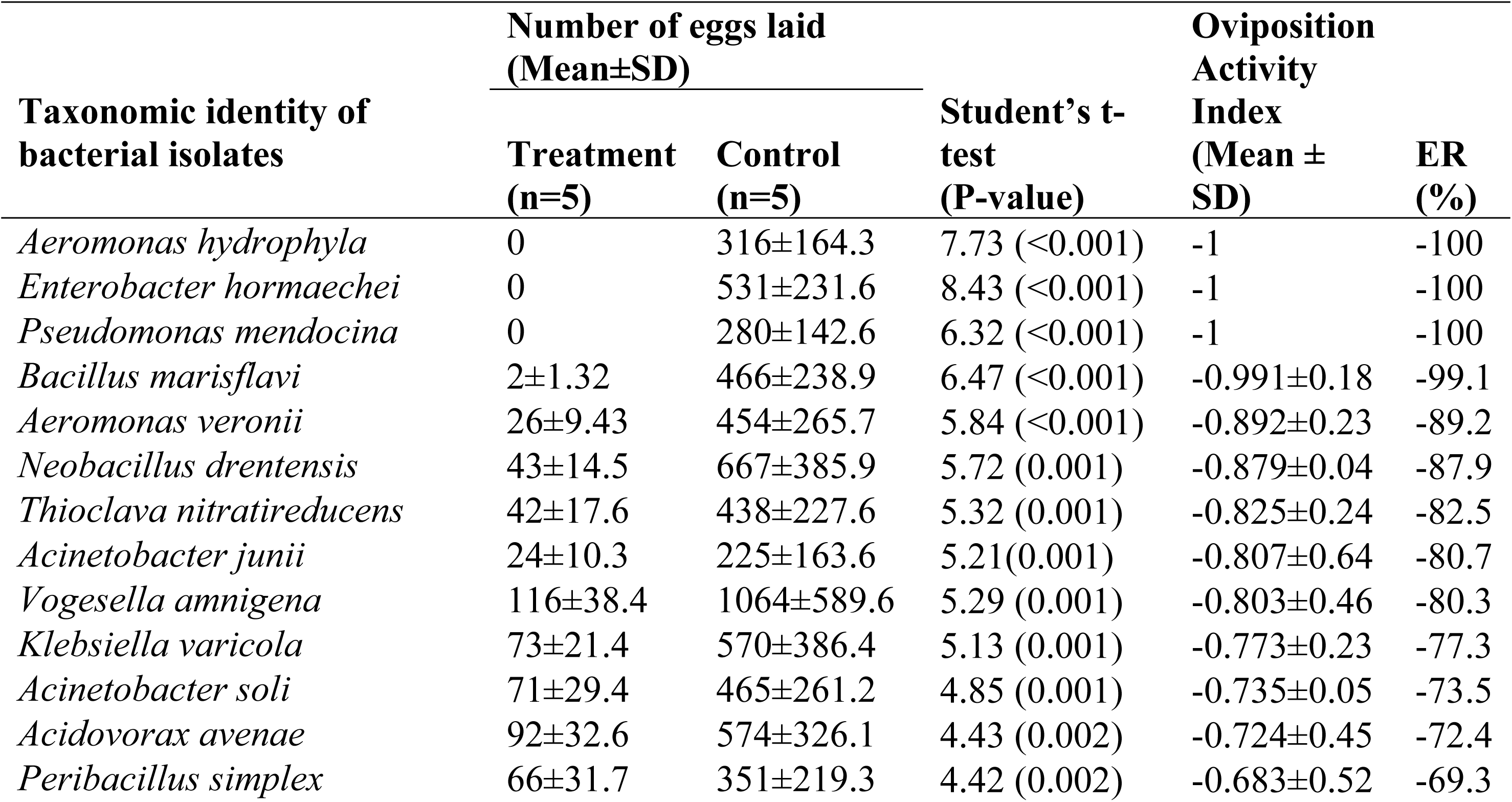
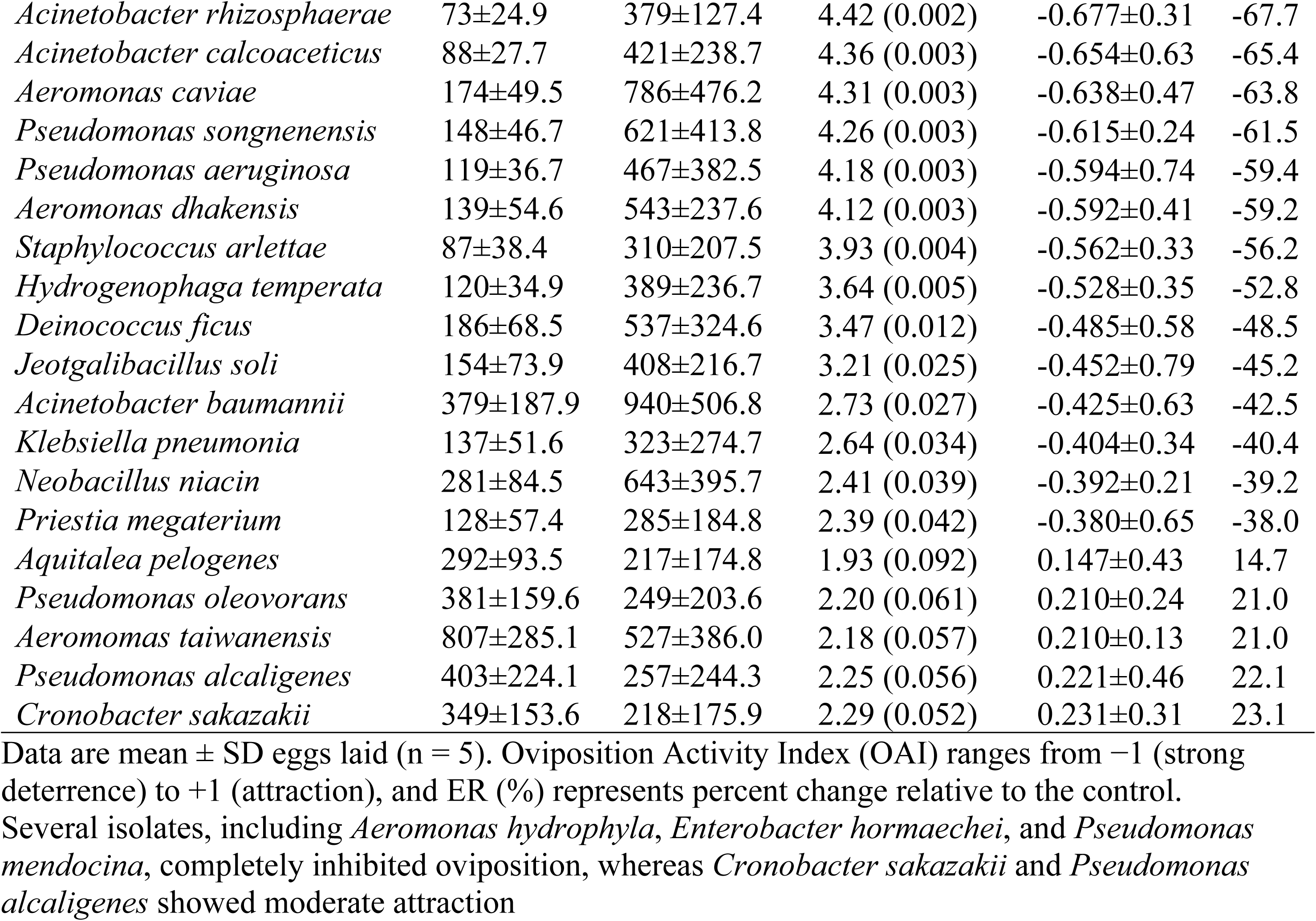
Oviposition responses of gravid mosquitoes to bacterial isolates from larval habitats.

Other isolates such as *Neobacillus drentensis*, *Thioclava nitratireducens*, *Acinetobacter junii*, and *Vogesella amnigena* also significantly suppressed oviposition by more than 80% relative to controls (P ≤ 0.001). A second group of isolates, including *Klebsiella varicola*, *Acinetobacter soli*, *Acidovorax avenae*, *Peribacillus simplex*, *Acinetobacter rhizosphaerae*, and *Acinetobacter calcoaceticus*, exhibited strong but comparatively moderate deterrent effects, reducing egg laying by 65–77% (OAI = −0.65 to −0.77; P ≤ 0.003). In contrast, isolates such as *Deinococcus ficus*, *Jeotgalibacillus soli*, *Acinetobacter baumannii*, *Klebsiella pneumoniae*, *Neobacillus niacin*, and *Priestia megaterium* displayed weaker but still significant deterrence, with egg reductions ranging from 38–49% (P ≤ 0.042). Notably, a small subset of isolates showed a trend towards oviposition attraction rather than deterrence. *Aquitalea pelogenes*, *Pseudomonas oleovorans*, *Aeromonas taiwanensis*, *Pseudomonas alcaligenes*, and *Cronobacter sakazakii* elicited higher egg deposition in treatment cups than in controls, although these differences were not statistically significant (P = 0.052–0.092), yielding positive OAI values (0.15–0.23) and positively suggesting weak attractant activity. Collectively, these findings demonstrate that bacterial communities associated with mosquito larval habitats strongly influence oviposition behavior, with most isolates acting as potent repellents, while a limited number may function as weak attractants.

### Analysis of Volatile Organic Compounds Associated with Bacterial Isolates

GC-MS analysis of volatiles emitted by the bacterial isolates revealed distinct chemical profiles for each species, with both qualitative and quantitative differences (Supplementary tables 2–11). *Neobacillus drentensis* produced a diverse array of compounds, dominated by 2,3,5,6- tetramethylphenol (15.3% of total peak area; 173.98 ng/µl), 1-(2,4,5-trimethylphenyl)ethanone (13.5%; 153.69 ng/µl), and 4′-hydroxyacetophenone (12.3%; 139.52 ng/µl), alongside several benzoic acid derivatives and phenolic ketones. In contrast, *Aeromonas hydrophila* emitted a simpler profile, with 3,5-diethylphenol representing the largest proportion (35.2%; 109.45 ng/µl), followed by m-ethyl acetophenone (26.0%; 80.79 ng/µl), indicating a prevalence of alkylphenols and acetophenone derivatives. *Thioclava nitratireducens*, although producing fewer compounds, released o-ethyl acetophenone as the major volatile (51.3%; 6.38 ng/µl), complemented by hydroxyacetophenone and substituted phenols. The volatile blend from *Pseudomonas mendocina* was more complex, with 4-vinylbenzoic acid as the most abundant compound (38.6%; 936.22 ng/µl), followed by 2-(1-methylethyl)benzoic acid (16.1%; 787.13 ng/µl) and 2,4,5- trimethylbenzaldehyde (9.0%; 711.54 ng/µl), highlighting the dominance of aromatic carboxylic acids and aldehydes. Similarly, *Acinetobacter junii* and *Rosellomeria marisflavi* produced acetophenone derivatives and substituted phenols as major constituents, with peak areas exceeding 50% in some cases. *Enterobacter hormaechei* was characterized by high emission of 1,1′-(1,4-phenylene)bis-ethanone (58.2%; 234.59 ng/µl) and 3,4,5-trimethylphenol (34.3%; 138.46 ng/µl), whereas *Aeromonas veronii* predominantly released 1-(2,3,4-trimethylphenyl)- ethanone (50.0%; 161.01 ng/µl) and 1-(2,4,5-trimethylphenyl)-ethanone (29.9%; 96.20 ng/µl).

*Volgesella amnigena* and *Acinetobacter calcoaceticus* showed more limited volatile spectra, but compounds such as 4-(1,1-dimethylethyl)-2-methylphenol (80.2%; 2.22 ng/µl) and 2,6-epidioxy-5-ethyl-3-iminomethyl-4-methyl-pyridine (30.3%; 1.85 ng/µl) were dominant, respectively. Across isolates, aromatic compounds, particularly alkylphenols, acetophenone derivatives, and benzoic acid derivatives, constituted the major chemical classes, suggesting conserved biosynthetic pathways among mosquito-associated bacteria. These quantitative and qualitative differences in VOC emission are likely to influence oviposition site selection by gravid *Anopheles* mosquitoes (Figure 2).

**Figure 2.**
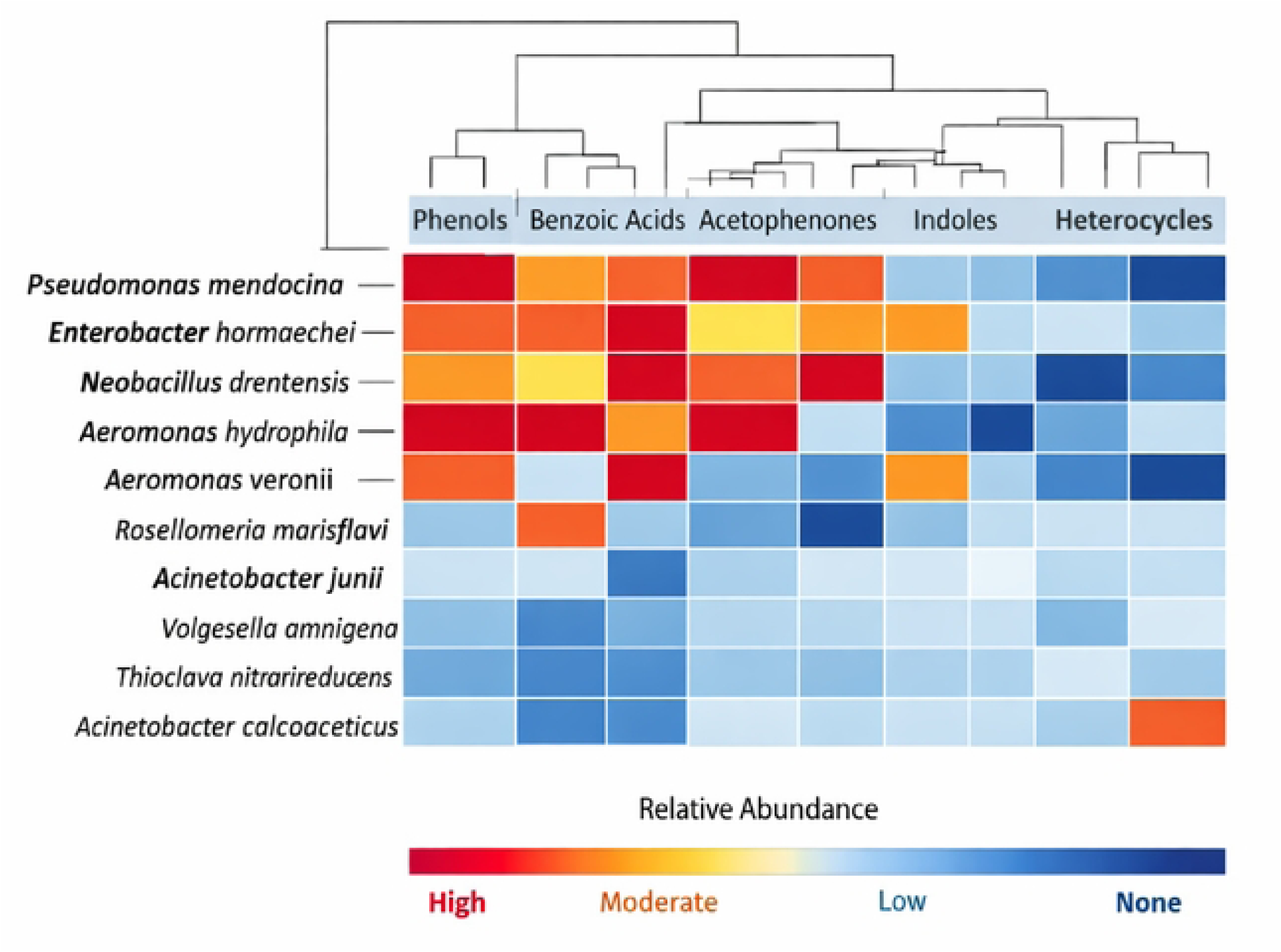
Volatile Profiles of Ten Bacterial Isolates Analyzed by GC-MS. The **rows** represent bacterial species. The **columns** represent chemical classes (Phenols, Benzoic Acids, Acetophenones, Indoles, Heterocycles).The **color gradient** (red → orange → light blue → dark blue) shows relative abundance, with red indicating high production and dark blue indicating absence. **Dendrograms** on both axes cluster species and chemical classes, revealing metabolic similarities (e.g., Pseudomonas mendocina and Enterobacter hormaechei cluster together due to strong benzoic acid and phenol output).

**Figure 3.**
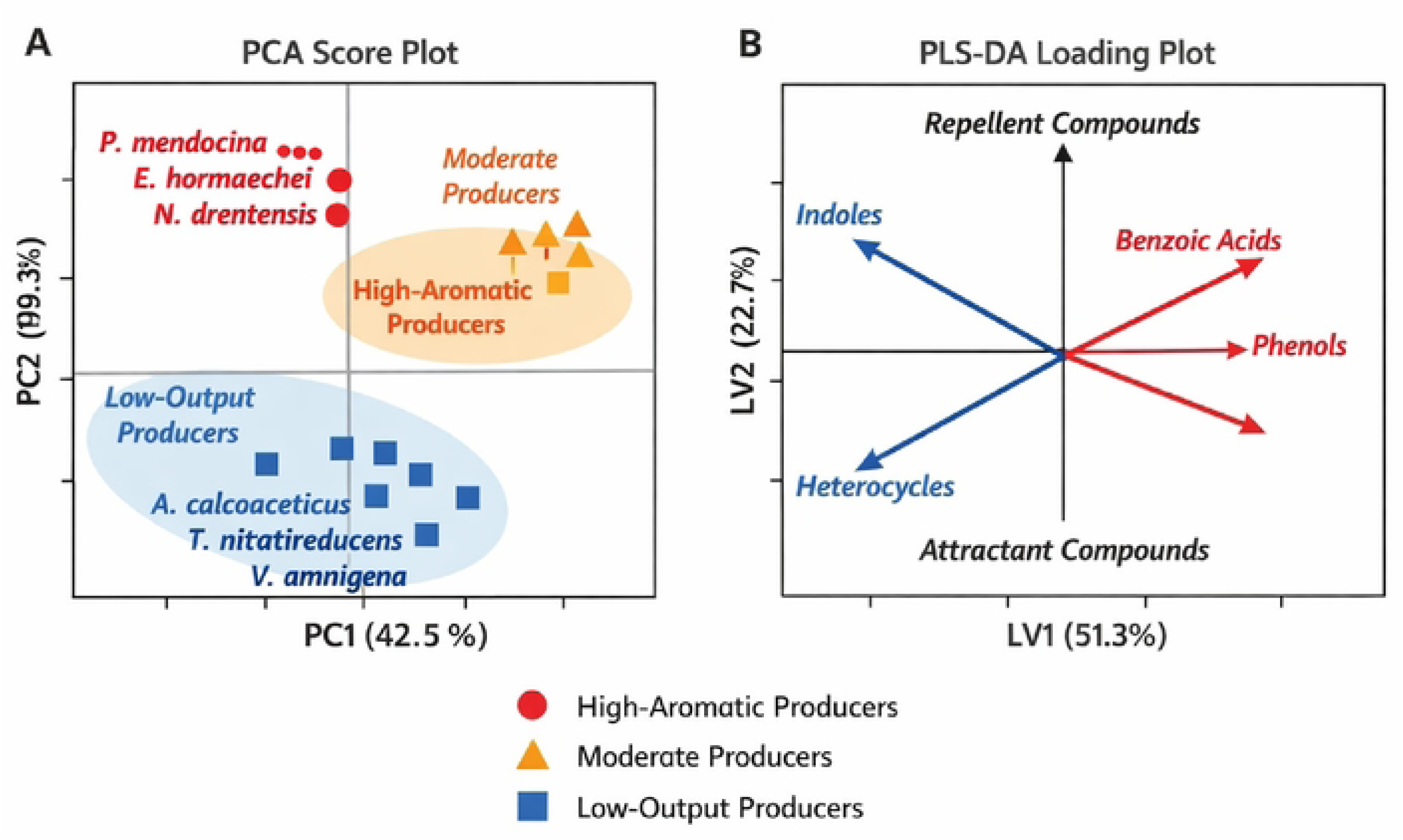
Multivariate analysis of volatile profiles from bacterial isolates associated with mosquito larval habitats. (A) **Principal Component Analysis (PCA) score plot** showing clustering of bacterial species based on volatile composition. Three distinct groups are evident: high-aromatic producers (Pseudomonas mendocina, Enterobacter hormaechei, Neobacillus drentensis), moderate producers (Aeromonas hydrophila, Aeromonas veronii, Rosellomeria marisflavi), and low-output producers (Acinetobacter calcoaceticus, Thioclava nitratireducens, Volgesella amnigena). PC1 (42.5 %) captures variance driven by phenols, benzoic acids, and acetophenones, while PC2 (19.3 %) separates isolates emitting heterocycles and indoles. **(B) Partial Least Squares Discriminant Analysis (PLS-DA) loading plot** illustrating the contribution of volatile classes to repellency or attraction. Positive loadings (LV1 > 0) correspond to benzoic acids and phenols, which are associated with strong repellency, whereas negative loadings (LV1 < 0) correspond to indoles and heterocycles, linked to weak attraction or neutral responses.

### Volatile chemical group composition of bacterial isolates

GC–MS analysis revealed that the volatile blends emitted by the bacterial isolates were dominated by benzenoid-derived compounds belonging to a limited number of major chemical groups (Table 2). Aromatic ketones, primarily acetophenone and propiophenone derivatives, constituted the largest fraction of detected compounds (41.8%), with trimethylacetophenones, hydroxyacetophenones, and diaryl ketones occurring as dominant components across multiple isolates. Phenolic compounds, mainly alkyl- and methoxy-substituted phenols, represented 19.6% of the total volatile composition, including tetramethylphenol, diethylphenol, and propofol. Aromatic carboxylic acids and their esters, largely benzoic acid and alkylbenzoic acid derivatives, accounted for a further 17.3%, with 4-vinylbenzoic acid and ethylbenzoic acid being particularly abundant in *Pseudomonas mendocina* and *Neobacillus drentensis*. Minor chemical groups included aromatic aldehydes (5.2%), aromatic ethers (4.6%), oxygenated heterocycles (4.0%), and nitrogen-containing heteroaromatics (4.2%), while aliphatic fatty acids and hydrocarbons contributed less than 3% of the total volatile blend. Overall, more than 80% of the detected volatiles belonged to aromatic functional classes typically associated with shikimate- and polyketide-related biosynthetic pathways, highlighting the predominance of benzenoid metabolism in these bacterial isolates and their potential role as semiochemically active compounds influencing mosquito oviposition behaviour.

**Table 2.** Summary of major chemical groups of volatile organic compounds emitted by bacterial isolates.

| Chemical Group | Representative Compounds | Number of Compounds | Relative abundance (%) |
| --- | --- | --- | --- |
| Aromatic ketones (acetophenones, propiophenones, diaryl ketones) | 1-(2,4,5-trimethylphenyl)ethanone, o-ethyl acetophenone, hydroxyacetophenone, diaryl ethanone | 24 | **41.8** |
| Phenols (alkylphenols, methoxyphenols, polyalkyl phenols) | 2,3,5,6-tetramethylphenol, 3,5-diethylphenol, propofol, eugenol, tert-butyl methylphenol | 18 | **19.6** |
| Aromatic carboxylic acids & esters (benzoic acid derivatives) | 4-vinylbenzoic acid, 2-ethylbenzoic acid, p-tert-butyl benzoic acid, cinnamic acid | 17 | **17.3** |
| Aromatic aldehydes | 2,4,5-trimethylbenzaldehyde, isophthalaldehyde, hydroxybenzaldehyde | 6 | **5.2** |
| Aromatic ethers & alkoxybenzenes | tert-butyl ethoxybenzene, anisaldehyde, methoxyphenols | 6 | **4.6** |
| Oxygenated heterocycles (chromanones, benzodioxins, coumarins) | Tetrahydromethylchromenone, benzodioxin acetophenone, acetylcoumarin | 5 | **4.0** |
| Nitrogen-containing aromatics & heterocycles | Indole, pyrimidinones, quinolinones, pyridines | 9 | **4.2** |
| Aliphatic fatty acids & hydrocarbons | Oleic acid, octadecanoic acid, pentadecanoic acid, heptacosane | 4 | **2.3** |
| Aromatic amides & nitriles | Ethylbenzamides, benzonitriles | 4 | **0.9** |
| **Total** |  | **93**<br>**compounds** | **100** |

**Table 2**. Major chemical groups of volatile organic compounds (VOCs) produced by bacterial isolates, with representative compounds, number of compounds, and relative abundance (%). These VOCs, including aromatic ketones, phenols, and carboxylic acid derivatives, potentially influence mosquito behavior and microbial interactions in breeding environments.

### Chemical Profile-Based Relationship Between Isolates

In the multivariate analysis of volatile profiles, Principal Component Analysis (PCA) revealed distinct clustering patterns among the bacterial isolates based on their chemical composition. The first two principal components accounted for over 60% of the total variance, effectively separating high-aromatic producers from moderate and low-output species. *Pseudomonas mendocina*, *Enterobacter hormaechei*, and *Neobacillus drentensis* grouped together along PC1, characterized by elevated levels of phenols, benzoic acids, and acetophenones. In contrast, *Aeromonas hydrophila*, *Aeromonas veronii*, and *Rosellomeria marisflavi* occupied an intermediate position, reflecting moderate production of ketones and phenols. Low-output isolates such as *Acinetobacter calcoaceticus*, *Thioclava nitratireducens*, and *Volgesella amnigena* clustered near the origin, indicating minimal volatile emission and a predominance of heterocyclic and indolic compounds. This separation underscores the metabolic diversity among larval habitat bacteria and their differential contributions to the volatile landscape.

Complementary Partial Least Squares Discriminant Analysis (PLS-DA) further clarified the chemical drivers of repellency and attraction. The first latent variable (LV1) distinguished isolates emitting high concentrations of benzoic acids and phenols, compounds strongly associated with oviposition deterrence, from those producing indoles and heterocycles, which correlated with weak attraction or neutral responses. The model achieved robust discrimination, confirming that aromatic abundance is the primary factor influencing mosquito avoidance.

Together, the PCA and PLS-DA results demonstrate a clear chemical polarity among isolates, where high-aromatic bacterial volatiles act as repellents, while low-level heterocyclic emissions may serve as subtle cues in oviposition site selection.

## DISCUSSION

### Isolation and Characterization of Bacteria

In the present study, we provide a lead to a potential link between larval habitat bacterial isolates and oviposition site selection by mosquito vectors. Pseudomonadota, Firmicutes, and Deinococcus in mosquito breeding sites play crucial ecological roles, which can have both positive and negative effects on the growth and development of mosquito larvae (16). Their ability to outcompete and suppress the growth of pathogenic bacteria in mosquito breeding sites prevents the mosquito larvae from being infected by these harmful bacteria (20). Additionally, these bacteria can effectively break down complex organic compounds into simpler forms, making them more accessible to mosquito larvae as a source of nutrition (11). On the contrary, some members of *Bacillus thuringiensis* produce toxins that can inhibit mosquito growth (30).

Furthermore, recent studies have suggested that the presence of bacteria in mosquito breeding sites can modulate the immune response of the mosquitoes (31,32).

Deinococcus species are known for their ability to survive in extreme environments, including high levels of radiation, desiccation, and low nutrient availability (33), and therefore their presence alone could prevent mosquito larvae from successfully proliferating. It is worth noting that the absence of mosquito larvae in a breeding site cannot be attributed solely to the presence of Deinococcus bacteria without further investigation. Although all the families of bacteria isolated have been reported in mosquito proliferation sites by other studies (18,34), their influence on mosquito breeding patterns remains unclear. While bacteria are known to be a source of food for the growth of mosquito larvae, they may also indicate the presence or absence of optimal conditions for larval growth in the proliferation site.

From this study, it is not clear why *Klebsiella* and *Bacillus* were mostly found in the negative sites, but it could be suggested that these bacteria impact negatively on the survival and development of mosquito larvae. Certain strains of *Bacillus* have been reported to produce toxins that can be lethal to mosquito larvae (30). Furthermore, if the larvae are exposed to these bacteria during their development, they might experience long-term effects that lead to reduced fertility or an impaired ability to lay viable eggs (35). It’s important to note that since *Klebsiella* and *Bacillus* can have negative effects on mosquito larvae, they might also play a role in biological control methods for mosquito populations. Some strains of these bacteria have been studied as potential biological control agents for mosquito populations due to their ability to suppress larval growth and development. Previous studies have demonstrated that bacterial communities present in mosquito proliferation sites vary based on factors such as mosquito species, type of habitat, and geographic location(36).

Various sources associated with mosquitoes, such as proliferation sites, mosquito midguts, and mosquito salivary glands, have been found to contain diverse types of bacteria (37). A wide range of bacterial groups, including *Aeromonas, Pseudomonas, Klebsiella, Staphylococcus, Bacillus, Acinetobacter,* and *Enterobacter* species, have also been isolated in the midguts of field-collected and laboratory-reared *Anopheles* mosquitoes (38). *Acinetobacter, Pseudomonas, Aeromonas, Aquitalea, Bacillus, Enterobacter, Klebsiella,* and *Staphylococcus* have been isolated from the larval habitats of *Anopheles darlingii* in Manaus (39).

The presence of *Aeromonas, Enterobacter,* and *Klebsiella* in water samples indicates that the water was contaminated with fecal waste (40). The high levels of these bacteria in the proliferation sites may affect the breeding patterns of *Anopheles gambiae,* which have been reported to proliferate in clean, unpolluted water in the environment (41). Investigations have also indicated a negative correlation between the number of *Anopheles* mosquito larvae and the population levels of fecal coliforms. These findings suggest that certain bacterial populations could potentially inhibit the growth of *Anopheles* larvae (42). It is possible that the presence of high levels of such bacteria in some potential breeding sites could explain why they did not have mosquito larvae. The altered chemical properties caused by these bacterial populations may have made the breeding sites unattractive to *Anopheles* mosquitoes, resulting in little to no mosquito larvae at those locations.

Dissolved oxygen in the water is more abundant in clean water than in polluted water, which is essential for mosquito larvae to survive (43). Additionally, clean water sources frequently harbor a diverse microbial community that can outcompete dangerous and pathogenic microorganisms, as well as a lower density of predatory organisms like other insect larvae, fish, or amphibians that feed on mosquito larvae (44). Clean water may emit specific chemical compounds that attract female mosquitoes, signaling the presence of suitable oviposition sites. Conversely, polluted water may produce different chemical cues that repel or deter the mosquitoes from laying their eggs.

Previous studies have also identified *Anopheles* mosquito larvae in unclean environmental water, particularly during the dry season (45). These findings are akin to those of the present study, underscoring the need for effective control measures to be employed during both the wet and dry seasons to prevent mosquito breeding and reduce the transmission of malaria. The presence of *Anopheles* mosquito larvae in some sites with pollution-associated bacteria could indicate that the mosquitoes can shift their preference for clean water to unclean water to survive through the dry season when proliferation sites are very scarce. The bacteria found in mosquito proliferation sites have been shown to have a significant impact on the diet and nutrition of mosquitoes, either as a direct food source or by breaking down large organic molecules into simpler compounds that can be more easily consumed by mosquito larvae (46).

Bacteria associated with mosquito larval habitats are predominantly located in the sediments and the rhizosphere of aquatic vegetation rather than in the water column itself, suggesting that chemical cues produced by these microorganisms may diffuse into the overlying water from these sources (47). The present study demonstrates that bacterial isolates from mosquito larval habitats exert strong and highly divergent effects on gravid mosquito oviposition behaviour.

These findings provide compelling evidence that habitat-associated bacteria are major chemical determinants of oviposition site selection in malaria vectors.

Several isolates, including *Aeromonas hydrophila*, *Enterobacter hormaechei*, and *Pseudomonas mendocina*, completely inhibited oviposition, with zero eggs recorded in all treatment replicates and oviposition activity indices (OAI) of –1.0. These results indicate extreme repellency and suggest that these taxa produce highly potent deterrent semiochemicals. The ecological relevance of such deterrence may reflect naturally occurring microbial succession dynamics in larval habitats, whereby gravid females avoid sites dominated by metabolically active or pathogenic bacteria that may reduce larval survival.

Strong but partial deterrence was observed for *Bacillus marisflavi*, *Aeromonas veronii*, *Neobacillus drentensis*, *Thioclava nitratireducens*, and *Acinetobacter junii*, all of which reduced egg deposition by more than 80% relative to controls. The consistently negative OAI values (– 0.80 to –0.99) indicate that these bacteria generate volatile profiles that are strongly unattractive to gravid females. Members of the genera *Aeromonas*, *Acinetobacter*, and *Bacillus* are well known producers of phenolic compounds, short-chain fatty acids, aromatic aldehydes, and heterocyclic metabolites, many of which function as oviposition deterrents at ecologically relevant concentrations. Intermediate levels of repellency were recorded for a diverse group of taxa, including *Klebsiella varicola*, *Acidovorax avenae*, *Peribacillus simplex*, *Acinetobacter rhizosphaerae*, *Aeromonas caviae*, and *Pseudomonas aeruginosa*, with egg reduction efficiencies ranging from 56% to 77%. These bacteria may produce complex volatile blends containing both attractant and deterrent components, resulting in net behavioural suppression.

In contrast, a small subset of isolates exhibited weak attraction or neutrality. *Aquitalea pelogenes*, *Pseudomonas oleovorans*, *Aeromonas taiwanensis*, *Pseudomonas alcaligenes*, and *Cronobacter sakazakii* displayed positive OAI values (0.15–0.23), although these responses were not statistically significant. These bacteria may produce volatile metabolites such as short-chain fatty acids, alcohols, or esters that mimic organic-rich breeding sites, thereby serving as oviposition cues. The observed behavioural responses likely reflect adaptive strategies evolved by mosquitoes to optimize offspring survival. Gravid females preferentially select habitats that provide adequate microbial food resources while avoiding environments dominated by pathogenic or toxic microbial communities. Consequently, bacterial composition functions as a reliable ecological indicator of habitat suitability, and mosquitoes exploit microbial semiochemicals as sensory proxies for larval fitness.

A study by Huang and others (48) reported that cultured bacteria originating from mosquito larval habitats had a negative effect on the oviposition behavior of *Anopheles gambiae,* suggesting that the mosquitoes are sensitive to bacterial-derived odors emanating from natural larval habitats and that some odors are repellent. It has been demonstrated by Eneh and others (49), that gravid *Anopheles gambiae* ss prefer to oviposit in tap water when the choice is a Bermuda grass hay infusion, and that all the volatile compounds identified and evaluated in Bermuda grass hay infusion were associated with bacterial activity. Lindh and others (50) reported that water from mosquito breeding sites contained bacteria, some of which elicited a positive oviposition response in gravid *Anopheles* mosquitoes; however, the results were inconclusive since no single attractive volatile compound was associated with any of the attractive bacteria, hence the source of attraction remained unknown. It may be interesting in the future to investigate the effect of a mixed culture of habitat-derived bacteria on oviposition behavior to elucidate potential synergistic or antagonistic effects.

The mosquitoes could have evolved the ability to detect and avoid such cues to seek optimal larval habitats that ensure the survival of their offspring based on previous negative experiences or genetic predisposition. It is suggested that these isolates carry out metabolic activities leading to the formation of metabolites that may influence the breeding and proliferation patterns of mosquitoes in the environment. Additionally, some bacteria produce toxins that act against mosquito larvae (51). Bacteria could alter water quality parameters or promote the growth of predators or parasites that could prey on mosquito larvae.

### Analysis of Volatile Organic Compounds Emitted by Bacteria

Ten bacterial species isolated from mosquito larval habitats were selected for characterization of their volatile metabolites through GC-MS due to their high repellence towards gravid *Anopheles gambiae* ss seeking oviposition sites. This is an indication that bacteria can emit diverse volatile compounds with different physicochemical characteristics and biological functions during their growth cycle, similar to what has been suggested in recent studies (52,53). Although microbial volatile organic compounds were previously considered secondary metabolites, recent findings have revealed that a majority of these compounds have significant biological functions (54).

These compounds are also believed to contribute significantly to intra- and inter-kingdom interactions where bacteria are involved (52). There is sufficient evidence suggesting that volatiles can diffuse to cover some considerable distance from their source, hence making them strategic semiochemicals for mediating both short- and long-range interactions between organisms (55).

Multivariate analysis of volatile emissions revealed clear chemical differentiation among bacterial isolates from mosquito larval habitats. The PCA score plot separated isolates into three clusters: high aromatic producers (*Pseudomonas mendocina*, *Enterobacter hormaechei*, *Neobacillus drentensis*), moderate producers (*Aeromonas hydrophila*, *Aeromonas veronii*, *Rosellomeria marisflavi*), and low-output producers (*Acinetobacter calcoaceticus*, *Thioclava nitratireducens*, *Volgesella amnigena*). Variance along PC1 (42.5%) was driven by phenols, benzoic acids, and acetophenones, compounds strongly linked to oviposition deterrence, while PC2 (19.3%) distinguished isolates emitting heterocycles and indoles, associated with weak attraction. The PLS-DA loading plot further clarified the ecological relevance of volatile classes. Positive loadings (LV1 > 0) corresponded to benzoic acids and phenols, reinforcing their role as repellents, whereas negative loadings (LV1 < 0) corresponded to indoles and heterocycles, linked to neutral or mildly attractive responses. This polarity highlights a chemical continuum in which aromatic abundance deters oviposition, while heterocyclic and indolic emissions may act as subtle cues of habitat suitability. Overall, these findings suggest that mosquito oviposition behavior is shaped by microbial volatile diversity rather than a single dominant compound.

High-aromatic isolates may serve as natural repellents for vector control, while low-output isolates producing indoles and heterocycles may inadvertently encourage oviposition.

Previous studies have reported that various bacterial groups emit distinct chemical signatures during their growth cycles, which are remarkably diverse in structure and chemical complexity, such that they can be used to differentiate between various groups of bacteria with high fidelity (56,57). This implies that there could be a complex interplay between the diversity, abundance, and distinctiveness of volatile chemical blends associated with these bacterial isolates, which modulate the oviposition site-seeking behavior of gravid *Anopheles gambiae* ss.

The volatile profiles of the bacterial isolates in this study were dominated by benzenoid-derived compounds, particularly aromatic ketones, phenols, and aromatic carboxylic acids, which together accounted for more than 80% of the detected components. These chemical groups are frequently implicated in the semiochemical mediation of mosquito oviposition behaviour. Early work demonstrated that bacteria-associated chemical cues significantly influence oviposition site selection by *Aedes aegypti*, with gravid females exhibiting preferences for water conditioned with specific bacterial isolates over sterile controls, indicating a role for bacterial volatiles in oviposition site attractiveness (58). Aromatic compounds such as phenols and carboxylic acids have been identified in hay infusion blends and bacterial headspace extracts that elicited electrophysiological and behavioural responses in *Culex* species, further supporting the behavioural relevance of these chemical groups (59). This indicates that the diversity and abundance of volatile organic compounds emitted by bacteria may vary within and between species due to changes in the culture medium coupled with growth conditions. Consequently, this may explain the huge diversity of volatiles emitted by the different bacteria. The production of volatile organic compounds by microorganisms is influenced by their genetic makeup, growth cycle, as well as the source and availability of nutrients. It is well known that the appearance of a recognizable volatile chemical profile in a bacterial species is largely attributable to its specific metabolic pathways (60).

Understanding the mechanisms behind gravid *Anopheles* mosquitoes’ avoidance behavior towards bacterial-infested water can have practical implications for malaria control strategies. By further investigating the specific chemical cues or microbial interactions that trigger avoidance, researchers can develop novel approaches to manipulating these factors in mosquito breeding sites. Targeting larval habitats by altering the microbial composition or using naturally occurring bacteria that are detrimental to *Anopheles* larvae could help reduce the mosquito population and interrupt malaria transmission. Alternatively, the identification and utilization of bacterial strains that promote the breeding of mosquito predators or parasites could present innovative biological control methods. By manipulating bacterial flora in mosquito breeding sites, it may be possible to disrupt the reproductive success of *Anopheles* mosquitoes and subsequently reduce malaria transmission rates on a larger scale. Identification and synthesis of the active volatile compounds could provide environmentally sustainable alternatives to conventional chemical control methods.

## Conclusion

The results highlight the central role of bacterial communities in shaping mosquito breeding ecology. Dominant genera, including *Aeromonas*, *Acinetobacter*, *Pseudomonas*, and *Bacillus*, were shown to influence the attractiveness or repellency of aquatic habitats. This underscores the importance of microbial composition as an ecological determinant of mosquito distribution, abundance, and habitat selection. The pronounced negative oviposition responses observed for several bacterial isolates suggest strong potential for application in vector control strategies.

Exploiting bacteria that deter oviposition could support the development of innovative interventions such as microbial-based larvicides, oviposition deterrents, or attract-and-kill systems. Furthermore, identification of the specific metabolites responsible for these effects may enable the development of synthetic analogues for environmentally sustainable mosquito control.

Notably, isolates such as *Pseudomonas mendocina*, *Neobacillus drentensis*, *Enterobacter hormaechei*, *Rosellomeria marisflavi*, *Aeromonas veronii*, and *Aeromonas hydrophila* produced distinct volatile organic compound (VOC) profiles associated with strong oviposition deterrence. These findings support the hypothesis that microbial-derived semiochemicals serve as critical cues in oviposition site selection, enabling mosquitoes to assess habitat suitability. Overall, this study provides new insights into the role of bacteria and their volatile metabolites in modulating oviposition behavior in *Anopheles gambiae s.s.*. It highlights the potential of leveraging microbe-mediated chemical ecology in the design of targeted, sustainable vector control strategies.

Further research should focus on isolating and characterizing the specific bioactive compounds involved and evaluating their applicability under field conditions.

## Acknowledgments

The authors wish to thank Festus Yaah, Mathews Wafula, and Joseph Khamisi for their support during the sample collection and laboratory experiments.

## CRediT Author Contributions statement

Josphat Mutinda: Writing – Original draft, Conceptualization, Data curation, Formal analysis, Funding acquisition, Investigation, Methodology, Project administration, Resources, Visualization. Kennedy O. Oduor: Writing – Review and editing, Data curation, Formal analysis, Investigation, Methodology, Resources, Visualization. Sammuel M. Mwamburi: Writing – Review and editing, Data curation, Formal analysis, Investigation, Methodology. Maurice O. Omolo: Writing – Review and editing, Conceptualization, Data curation, Formal analysis, Investigation, Supervision, Validation. Regina M. Ntabo: Writing – Review and editing, Conceptualization, Formal analysis, Investigation, Methodology, Supervision. James M. Gathiru: Writing – Original draft, Conceptualization, Data curation, Formal analysis, Investigation.

Joseph Mwangangi: Writing – Review and editing, Conceptualization, Formal analysis, Methodology, Supervision, Validation. James Nonoh: Writing – Review and editing, Conceptualization, Data curation, Formal analysis, Funding acquisition, Investigation,

Methodology, Supervision, Validation. All authors have read and approved the final version of the manuscript and agree to be accountable for all aspects of the work, including ensuring the accuracy and integrity of the study.

## Data availability statement

All data generated or analysed during this study are included in this published article and its supplementary information file. The 16S rRNA gene sequences generated in this study have been deposited in the National Center for Biotechnology Information (NCBI) GenBank database under the assession numbers PX869225 – PX869248 (GenBank Submission SUB15935019)

## Declaration of Generative AI Use

The authors acknowledge the use of generative AI, ChatGPT and OpenAI in the preparation of this manuscript for purposes of language refinement, formatting, and organization of content. The AI tools were not used to generate scientific ideas, experimental data, or conclusions. All outputs were carefully reviewed and edited by the authors to ensure accuracy and compliance with scientific standards. The authors accept full responsibility for the final content of the manuscript.

## References

1. Takken W, Verhulst NO. Host preferences of blood-feeding mosquitoes. Annu Rev Entomol. 2013;58:433–53. doi:10.1146/annurev-ento-120811-153618 PubMed PMID: 23020619.

2. Owusu-Ofori A, Gadzo D, Bates I. Transfusion-transmitted malaria: donor prevalence of parasitaemia and a survey of healthcare workers knowledge and practices in a district hospital in Ghana. Malar J. 2016 Apr 23;15:234. doi:10.1186/s12936-016-1289-3 PubMed PMID: 27108087; PubMed Central PMCID: PMC4842271.

3. O’Meara WP, Bejon P, Mwangi TW, Okiro EA, Peshu N, Snow RW, et al. Effect of a fall in malaria transmission on morbidity and mortality in Kilifi, Kenya. Lancet. 2008 Nov 1;372(9649):1555–62. doi:10.1016/S0140-6736(08)61655-4 PubMed PMID: 18984188; PubMed Central PMCID: PMC2607008.

4. Snow RW, Kibuchi E, Karuri SW, Sang G, Gitonga CW, Mwandawiro C, et al. Changing Malaria Prevalence on the Kenyan Coast since 1974: Climate, Drugs and Vector Control. PLOS ONE. 2015 Jun 24;10(6):e0128792. doi:10.1371/journal.pone.0128792

5. Chirebvu E, Chimbari MJ. Characteristics of Anopheles arabiensis larval habitats in Tubu village, Botswana. J Vector Ecol J Soc Vector Ecol. 2015 Jun;40(1):129–38. doi:10.1111/jvec.12141 PubMed PMID: 26047193.

6. Getachew D, Balkew M, Tekie H. Anopheles larval species composition and characterization of breeding habitats in two localities in the Ghibe River Basin, southwestern Ethiopia. Malar J. 2020 Feb 11;19:65. doi:10.1186/s12936-020-3145-8 PubMed PMID: 32046734; PubMed Central PMCID: PMC7014609.

7. Pluess B, Tanser FC, Lengeler C, Sharp BL. Indoor residual spraying for preventing malaria. Cochrane Database Syst Rev. 2010 Apr 14;2010(4):CD006657. doi:10.1002/14651858.CD006657.pub2 PubMed PMID: 20393950; PubMed Central PMCID: PMC6532743.

8. Ferguson HM, Dornhaus A, Beeche A, Borgemeister C, Gottlieb M, Mulla MS, et al. Ecology: A Prerequisite for Malaria Elimination and Eradication. PLOS Med. 2010 Aug 3;7(8):e1000303. doi:10.1371/journal.pmed.1000303

9. Kiuru CW, Oyieke FA, Mukabana WR, Mwangangi J, Kamau L, Muhia-Matoke D. Status of insecticide resistance in malaria vectors in Kwale County, Coastal Kenya. Malar J. 2018 Jan 5;17(1):3. doi:10.1186/s12936-017-2156-6

10. Fillinger U, Lindsay SW. Larval source management for malaria control in Africa: myths and reality. Malar J. 2011 Dec 13;10(1):353. doi:10.1186/1475-2875-10-353

11. Souza RS, Virginio F, Riback TIS, Suesdek L, Barufi JB, Genta FA. Microorganism-Based Larval Diets Affect Mosquito Development, Size and Nutritional Reserves in the Yellow Fever Mosquito Aedes aegypti (Diptera: Culicidae). Front Physiol. 2019 Apr 9;10. doi:10.3389/fphys.2019.00152

12. Majambere S, Fillinger U, Sayer DR, Green C, Lindsay SW. Spatial distribution of mosquito larvae and the potential for targeted larval control in The Gambia. Am J Trop Med Hyg. 2008 Jul;79(1):19–27. PubMed PMID: 18606759.

13. Himeidan YE, Temu EA, El Rayah EA, Munga S, Kweka EJ. Chemical Cues for Malaria Vectors Oviposition Site Selection: Challenges and Opportunities. J Insects. 2013;2013(1):685182. doi:10.1155/2013/685182

14. Sumba LA, Guda TO, Deng AL, Hassanali A, Beier JC, Knols BGJ. Mediation of oviposition site selection in the African malaria mosquito Anopheles gambiae (Diptera: Culicidae) by semiochemicals of microbial origin. Int J Trop Insect Sci. 2004 Sep 1;24:260–5. doi:10.1079/IJT200433

15. Wang Y, Iii TMG, Kukutla P, Yan G, Xu J. Dynamic Gut Microbiome across Life History of the Malaria Mosquito Anopheles gambiae in Kenya. PLOS ONE. 2011 Sep 21;6(9):e24767. doi:10.1371/journal.pone.0024767

16. Ranasinghe HAK, Amarasinghe LD. Naturally Occurring Microbiota Associated with Mosquito Breeding Habitats and Their Effects on Mosquito Larvae. BioMed Res Int. 2020 Dec 14;2020:4065315. doi:10.1155/2020/4065315 PubMed PMID: 33381553; PubMed Central PMCID: PMC7755482.

17. Poonam S, Paily KP, Balaraman K. Oviposition attractancy of bacterial culture filtrates: response of Culex quinquefasciatus. Mem Inst Oswaldo Cruz. 2002 Apr;97(3):359–62. doi:10.1590/s0074-02762002000300015 PubMed PMID: 12048566.

18. Chukalo* E, Abate D. Bacterial populations of mosquito breeding habitats in relation to maize pollen in Asendabo, south western Ethiopia. Afr J Microbiol Res. 2017 Jan 14;11(2):55–64. doi:10.5897/AJMR2016.8287

19. Omoke D, Kipsum M, Otieno S, Esalimba E, Sheth M, Lenhart A, et al. Western Kenyan Anopheles gambiae showing intense permethrin resistance harbour distinct microbiota. Malar J. 2021 Feb 8;20:77. doi:10.1186/s12936-021-03606-4 PubMed PMID: 33557825; PubMed Central PMCID: PMC7869237.

20. Girard M, Martin E, Vallon L, Raquin V, Bellet C, Rozier Y, et al. Microorganisms Associated with Mosquito Oviposition Sites: Implications for Habitat Selection and Insect Life Histories. Microorganisms. 2021 Aug;9(8):1589. doi:10.3390/microorganisms9081589

21. Minas K, McEwan NR, Newbold CJ, Scott KP. Optimization of a high-throughput CTAB- based protocol for the extraction of qPCR-grade DNA from rumen fluid, plant and bacterial pure cultures. FEMS Microbiol Lett. 2011 Dec;325(2):162–9. doi:10.1111/j.1574-6968.2011.02424.x PubMed PMID: 22029887.

22. Costello EK, Lauber CL, Hamady M, Fierer N, Gordon JI, Knight R. Bacterial Community Variation in Human Body Habitats Across Space and Time. Science. 2009 Dec 18;326(5960):1694–7. doi:10.1126/science.1177486 PubMed PMID: 19892944; PubMed Central PMCID: PMC3602444.

23. Lee PY, Costumbrado J, Hsu CY, Kim YH. Agarose gel electrophoresis for the separation of DNA fragments. J Vis Exp JoVE. 2012 Apr 20;(62):3923. doi:10.3791/3923 PubMed PMID: 22546956; PubMed Central PMCID: PMC4846332.

24. Desjardins PR, Conklin DS. Microvolume quantitation of nucleic acids. Curr Protoc Mol Biol. 2011 Jan;Appendix 3:3J. doi:10.1002/0471142727.mba03js93 PubMed PMID: 21225636.

25. Herrera-Varela M, Lindh J, Lindsay SW, Fillinger U. Habitat discrimination by gravid Anopheles gambiae sensu lato – a push-pull system. Malar J. 2014 Apr 2;13:133. doi:10.1186/1475-2875-13-133 PubMed PMID: 24693951; PubMed Central PMCID: PMC3975139.

26. Okal MN, Lindh JM, Torr SJ, Masinde E, Orindi B, Lindsay SW, et al. Analysing the oviposition behaviour of malaria mosquitoes: design considerations for improving two- choice egg count experiments. Malar J. 2015 Jun 20;14:250. doi:10.1186/s12936-015-0768-2 PubMed PMID: 26088669; PubMed Central PMCID: PMC4474426.

27. Wang P, Sun M, Ren J, Djuric Z, Fisher GJ, Wang X, et al. Gas chromatography-mass spectrometry analysis of effects of dietary fish oil on total fatty acid composition in mouse skin. Sci Rep. 2017 Feb 14;7:42641. doi:10.1038/srep42641 PubMed PMID: 28195161; PubMed Central PMCID: PMC5307384.

28. Guo B, Yuan Y. A comparative review of methods for comparing means using partially paired data. Stat Methods Med Res. 2017 Jun;26(3):1323–40. doi:10.1177/0962280215577111 PubMed PMID: 25834090.

29. Kramer WL, Mulla MS. Oviposition Attractants and Repellents of Mosquitoes: Oviposition Responses ofCulex1Mosquitoes to Organic Infusions2. Environ Entomol. 1979 Dec 1;8(6):1111–7. doi:10.1093/ee/8.6.1111

30. Silva-Filha MHNL, Romão TP, Rezende TMT, Carvalho K da S, Gouveia de Menezes HS, Alexandre do Nascimento N, et al. Bacterial Toxins Active against Mosquitoes: Mode of Action and Resistance. Toxins. 2021 Aug;13(8):523. doi:10.3390/toxins13080523

31. Djadid ND, Jazayeri H, Raz A, Favia G, Ricci I, Zakeri S. Identification of the Midgut Microbiota of An. stephensi and An. maculipennis for Their Application as a Paratransgenic Tool against Malaria. PLOS ONE. 2011 Dec 6;6(12):e28484. doi:10.1371/journal.pone.0028484

32. Engel P, Moran NA. The gut microbiota of insects – diversity in structure and function. FEMS Microbiol Rev. 2013 Sep 1;37(5):699–735. doi:10.1111/1574-6976.12025

33. Zeng YH, Shen FT, Tan CC, Huang CC, Young CC. The flexibility of UV-inducible mutation in Deinococcus ficus as evidenced by the existence of the imuB-dnaE2 gene cassette and generation of superior feather degrading bacteria. Microbiol Res. 2011 Dec 20;167(1):40–7. doi:10.1016/j.micres.2011.02.008 PubMed PMID: 21459566.

34. Mosquera KD, Martinez Villegas LE, Pidot SJ, Sharif C, Klimpel S, Stinear TP, et al. Multi- Omic Analysis of Symbiotic Bacteria Associated With Aedes aegypti Breeding Sites. Front Microbiol. 2021 Aug 12;12. doi:10.3389/fmicb.2021.703711

35. Dickson LB, Jiolle D, Minard G, Moltini-Conclois I, Volant S, Ghozlane A, et al. Carryover effects of larval exposure to different environmental bacteria drive adult trait variation in a mosquito vector. Sci Adv. 2017 Aug;3(8):e1700585. doi:10.1126/sciadv.1700585 PubMed PMID: 28835919; PubMed Central PMCID: PMC5559213.

36. Caragata EP, Otero LM, Tikhe CV, Barrera R, Dimopoulos G. Microbial Diversity of Adult Aedes aegypti and Water Collected from Different Mosquito Aquatic Habitats in Puerto Rico. Microb Ecol. 2022 Jan;83(1):182–201. doi:10.1007/s00248-021-01743-6 PubMed PMID: 33860847; PubMed Central PMCID: PMC11328149.

37. Gabrieli P, Caccia S, Varotto-Boccazzi I, Arnoldi I, Barbieri G, Comandatore F, et al. Mosquito Trilogy: Microbiota, Immunity and Pathogens, and Their Implications for the Control of Disease Transmission. Front Microbiol. 2021 Apr 6;12. doi:10.3389/fmicb.2021.630438

38. Tuanudom R, Yurayart N, Rodkhum C, Tiawsirisup S. Diversity of midgut microbiota in laboratory-colonized and field-collected Aedes albopictus (Diptera: Culicidae): A preliminary study. Heliyon. 2021 Oct;7(10):e08259. doi:10.1016/j.heliyon.2021.e08259 PubMed PMID: 34765765; PubMed Central PMCID: PMC8569434.

39. Rocha EM, Marinotti O, Serrão DM, Correa LV, Katak R de M, de Oliveira JC, et al. Culturable bacteria associated with Anopheles darlingi and their paratransgenesis potential. Malar J. 2021 Jan 13;20(1):40. doi:10.1186/s12936-020-03574-1

40. ªahin S, Sivri N, Akpinar I, Çinçin ZB, Sönmez VZ. A comprehensive bibliometric overview: antibiotic resistance and Escherichia coli in natural water. Environ Sci Pollut Res Int. 2021 May 6;28(25):32256–63. doi:10.1007/s11356-021-14084-1 PubMed PMID: 33959839; PubMed Central PMCID: PMC8102055.

41. Emidi B, Kisinza WN, Mmbando BP, Malima R, Mosha FW. Effect of physicochemical parameters on Anopheles and Culex mosquito larvae abundance in different breeding sites in a rural setting of Muheza, Tanzania. Parasit Vectors. 2017 Jun 24;10(1):304. doi:10.1186/s13071-017-2238-x PubMed PMID: 28645303; PubMed Central PMCID: PMC5482952.

42. Nilsson LKJ, Sharma A, Bhatnagar RK, Bertilsson S, Terenius O. Presence of Aedes and Anopheles mosquito larvae is correlated to bacteria found in domestic water-storage containers. FEMS Microbiol Ecol. 2018 Jun 1;94(6). doi:10.1093/femsec/fiy058 PubMed PMID: 29617987.

43. Mamai W, Lees RS, Maiga H, Gilles JRL. Reusing larval rearing water and its effect on development and quality of Anopheles arabiensis mosquitoes. Malar J. 2016 Mar 16;15:169. doi:10.1186/s12936-016-1227-4 PubMed PMID: 26984183; PubMed Central PMCID: PMC4793705.

44. Dida GO, Gelder FB, Anyona DN, Abuom PO, Onyuka JO, Matano AS, et al. Presence and distribution of mosquito larvae predators and factors influencing their abundance along the Mara River, Kenya and Tanzania. SpringerPlus. 2015 Mar 20;4(1):136. doi:10.1186/s40064-015-0905-y

45. Gimnig JE, Ombok M, Kamau L, Hawley WA. Characteristics of larval anopheline (Diptera: Culicidae) habitats in Western Kenya. J Med Entomol. 2001 Mar;38(2):282–8. doi:10.1603/0022-2585-38.2.282 PubMed PMID: 11296836.

46. Tchioffo MT, Boissière A, Churcher TS, Abate L, Gimonneau G, Nsango SE, et al. Modulation of malaria infection in Anopheles gambiae mosquitoes exposed to natural midgut bacteria. PloS One. 2013;8(12):e81663. doi:10.1371/journal.pone.0081663 PubMed PMID: 24324714; PubMed Central PMCID: PMC3855763.

47. Orondo PW, Ochwedo KO, Atieli H, Yan G, Githeko AK, Nyanjom SG. Effects of bacterial composition and aquatic habitat metabolites on malaria vector larval availability in irrigated and non-irrigated sites of Homa Bay county, western Kenya. PLOS ONE. 2023 Jun 2;18(6):e0286509. doi:10.1371/journal.pone.0286509

48. Huang J, Walker ED, Otienoburu PE, Amimo F, Vulule J, Miller JR. Laboratory tests of oviposition by the African malaria mosquito, Anopheles gambiae, on dark soil as influenced by presence or absence of vegetation. Malar J. 2006 Oct 12;5:88. doi:10.1186/1475-2875-5-88 PubMed PMID: 17038187; PubMed Central PMCID: PMC1618395.

49. Eneh LK, Fillinger U, Karlson AKB, Rajarao GK, Lindh J. Anopheles arabiensis oviposition site selection in response to habitat persistence and associated physicochemical parameters, bacteria and volatile profiles. Med Vet Entomol. 2019 Mar;33(1):56–67. doi:10.1111/mve.12336 PubMed PMID: 30168151; PubMed Central PMCID: PMC6359949.

50. Lindh JM, Kännaste A, Knols BGJ, Faye I, Borg-Karlson AK. Oviposition responses of Anopheles gambiae s.s. (Diptera: Culicidae) and identification of volatiles from bacteria- containing solutions. J Med Entomol. 2008 Nov;45(6):1039–49. doi:10.1603/0022-2585(2008)45%5B1039:oroags%5D2.0.co;2 PubMed PMID: 19058627.

51. Suryadi BF, Yanuwiadi B, Ardyati T, Suharjono null. Isolation of Bacillus sphaericus from Lombok Island, Indonesia, and Their Toxicity against Anopheles aconitus. Int J Microbiol. 2015;2015:854709. doi:10.1155/2015/854709 PubMed PMID: 26788061; PubMed Central PMCID: PMC4691609.

52. Schulz-Bohm K, Martín-Sánchez L, Garbeva P. Microbial Volatiles: Small Molecules with an Important Role in Intra- and Inter-Kingdom Interactions. Front Microbiol. 2017 Dec 12;8. doi:10.3389/fmicb.2017.02484

53. Weisskopf L, Schulz S, Garbeva P. Microbial volatile organic compounds in intra-kingdom and inter-kingdom interactions. Nat Rev Microbiol. 2021 Jun;19(6):391–404. doi:10.1038/s41579-020-00508-1 PubMed PMID: 33526910.

54. Schmidt R, Etalo DW, de Jager V, Gerards S, Zweers H, de Boer W, et al. Microbial Small Talk: Volatiles in Fungal–Bacterial Interactions. Front Microbiol. 2016 Jan 5;6. doi:10.3389/fmicb.2015.01495

55. Maffei ME, Gertsch J, Appendino G. Plant volatiles: production, function and pharmacology. Nat Prod Rep. 2011 Aug;28(8):1359–80. doi:10.1039/c1np00021g PubMed PMID: 21670801.

56. Chen J, Tang JN, Hu KL, Zhao YY, Tang C. The production characteristics of volatile organic compounds and their relation to growth status of *Staphylococcus aureus* in milk environment. J Dairy Sci. 2018 Jun 1;101(6):4983–91. doi:10.3168/jds.2017-13629

57. Rees CA, Nordick KV, Franchina FA, Lewis AE, Hirsch EB, Hill JE. Volatile metabolic diversity of Klebsiella pneumoniae in nutrient-replete conditions. Metabolomics Off J Metabolomic Soc. 2017 Feb;13(2):18. doi:10.1007/s11306-016-1161-z PubMed PMID: 30464740; PubMed Central PMCID: PMC6241307.

58. Ponnusamy L, Schal C, Wesson DM, Arellano C, Apperson CS. Oviposition responses of Aedes mosquitoes to bacterial isolates from attractive bamboo infusions. Parasit Vectors. 2015 Sep 23;8(1):486. doi:10.1186/s13071-015-1068-y

59. Wooding M, Naudé Y, Rohwer E, Bouwer M. Controlling mosquitoes with semiochemicals: a review. Parasit Vectors. 2020 Feb 17;13(1):80. doi:10.1186/s13071-020-3960-3

60. Cheng X, Cordovez V, Etalo DW, van der Voort M, Raaijmakers JM. Role of the GacS Sensor Kinase in the Regulation of Volatile Production by Plant Growth-Promoting Pseudomonas fluorescens SBW25. Front Plant Sci. 2016 Nov 18;7. doi:10.3389/fpls.2016.01706

